# SARS-CoV-2 accessory proteins ORF7a and ORF3a use distinct mechanisms to downregulate MHC-I surface expression

**DOI:** 10.1101/2022.05.17.492198

**Authors:** Najla Arshad, Maudry Laurent-Rolle, Wesam S Ahmed, Jack Chun-Chieh Hsu, Susan M Mitchell, Joanna Pawlak, Debrup Sengupta, Kabir H Biswas, Peter Cresswell

## Abstract

Major histocompatibility complex class I (MHC-I) molecules, which are dimers of a glycosylated polymorphic transmembrane heavy chain and the small protein β_2_-microglobulin (β_2_m), bind peptides in the endoplasmic reticulum that are generated by the cytosolic turnover of cellular proteins. In virus-infected cells these peptides may include those derived from viral proteins. Peptide-MHC-I complexes then traffic through the secretory pathway and are displayed at the cell surface where those containing viral peptides can be detected by CD8^+^ T lymphocytes that kill infected cells. Many viruses enhance their *in vivo* survival by encoding genes that downregulate MHC-I expression to avoid CD8^+^ T cell recognition. Here we report that two accessory proteins encoded by SARS-CoV-2, the causative agent of the ongoing COVID-19 pandemic, downregulate MHC-I expression using distinct mechanisms. One, ORF3a, a viroporin, reduces global trafficking of proteins, including MHC-I, through the secretory pathway. The second, ORF7a, interacts specifically with the MHC-I heavy chain, acting as a molecular mimic of β_2_m to inhibit its association. This slows the exit of properly assembled MHC-I molecules from the endoplasmic reticulum. We demonstrate that ORF7a reduces antigen presentation by the human MHC-I allele HLA-A*02:01. Thus, both ORF3a and ORF7a act post-translationally in the secretory pathway to lower surface MHC-I expression, with ORF7a exhibiting a novel and specific mechanism that allows immune evasion by SARS-CoV-2.

**Significance Statement:** Viruses may down-regulate MHC class I expression on infected cells to avoid elimination by cytotoxic T cells. We report that the accessory proteins ORF7a and ORF3a of SARS-CoV-2 mediate this function and delineate the two distinct mechanisms involved. While ORF3a inhibits global protein trafficking to the cell surface, ORF7a acts specifically on MHC-I by competing with β_2_m for binding to the MHC-I heavy chain. This is the first account of molecular mimicry of β_2_m as a viral mechanism of MHC-I down-regulation to facilitate immune evasion.

## Introduction

Major histocompatibility complex class I (MHC-I) molecules are key players in mediating adaptive immune responses to viral infections. These molecules are heterodimers of a glycosylated transmembrane heavy chain and the small protein β_2_-microglobulin (β_2_m) that are synthesized and assembled in the endoplasmic reticulum (ER)(1). MHC-I is then loaded with peptides derived from the turnover of cellular proteins, including viral proteins in infected cells. Peptide loading is facilitated by the concerted action of proteins that comprise the peptide loading complex (PLC), namely the *t*ransporter associated with *a*ntigen *p*rocessing (TAP), which imports cellular and viral peptides from the cytosol into the ER, the peptide editor tapasin, and the ER chaperone proteins ERp57 and calreticulin(1). Peptide-MHC-I molecules then traffic to the surface of cells where self or virus-derived, non-self-peptides are exhibited to the immune system. This is the antigen processing and presentation pathway. Components of the adaptive arm of the immune system, particularly CD8^+^ T cell also known as cytotoxic T lymphocytes or CTLs, identify and kill cells that express non-self-peptides(1).

Viruses downregulate the expression of MHC-I by various mechanisms to avoid elimination by adaptive immune responses. This can involve transcriptional downregulation of MHC-I genes(2) or post-translational inhibition of MHC-I function(3, 4). In the latter case several steps in the antigen processing pathway are targeted by viruses. Examples include the herpes simplex virus ICP47 protein that inhibits TAP to prevent the import of cytosolic peptides, including virus-derived peptides, into the ER(3); the adenovirus E3/19K protein that prevents the interaction of MHC-I with the PLC(3) thereby interfering with effective peptide loading; and human papilloma virus E5 protein that retains MHC-I in the ER(5). Several viruses also encode pore-forming viral proteins or viroporins, which integrate into the host membranes, usually the ER, disrupting ion homeostasis(6) and inhibiting host protein trafficking through the secretory pathway. Examples include the influenza A virus M2 protein, poliovirus 2B and 3A proteins, coxsackievirus 2B protein, and SARS-CoV-1 ORF3a protein(7, 8). Cellular expression of poliovirus 3A viroporin(9) and coxsackievirus 2B viroporin(10) has been shown to reduce antigen presentation by MHC-I.

This study focuses on identifying SARS-CoV-2 gene products that post-translationally downregulate MHC-I expression and the mechanisms involved. SARS-CoV-2 is the causative agent of COVID-19, the ongoing pandemic that has afflicted the world for over two years, despite the availability of vaccines and emerging antiviral therapies (11). Thus, there remains a need to understand the molecular mechanisms of SARS-CoV-2 pathogenesis to aid the development of more effective, second generation anti-viral therapies. SARS-CoV-2 is a single, positive-stranded, enveloped RNA virus that encodes a number of translatable open reading frames (ORFs). These encode three main categories of proteins-non-structural, structural, and accessory proteins (Fig. 1*A*). Infection of cells with SARS-Cov-2 has been reported to downregulate MHC-I gene expression and reduce the surface expression of a specific MHC-I allele, HLA-A*02:01 (HLA-A2) (12, 13). Two accessory proteins have been implicated in this process. ORF6 suppressed the transcriptional upregulation of MHC-I observed upon viral infection (12) and expression of ORF8 post-translationally decreased overall levels of MHC-I. ORF8 was shown to interact with HLA-A2 and direct it to autophagosomes for degradation, thus depleting the intracellular pool and in turn lowering surface HLA-A2, leading to reduced recognition and killing of ORF8-expressing cells by CTLs (13).

**Figure 1.**
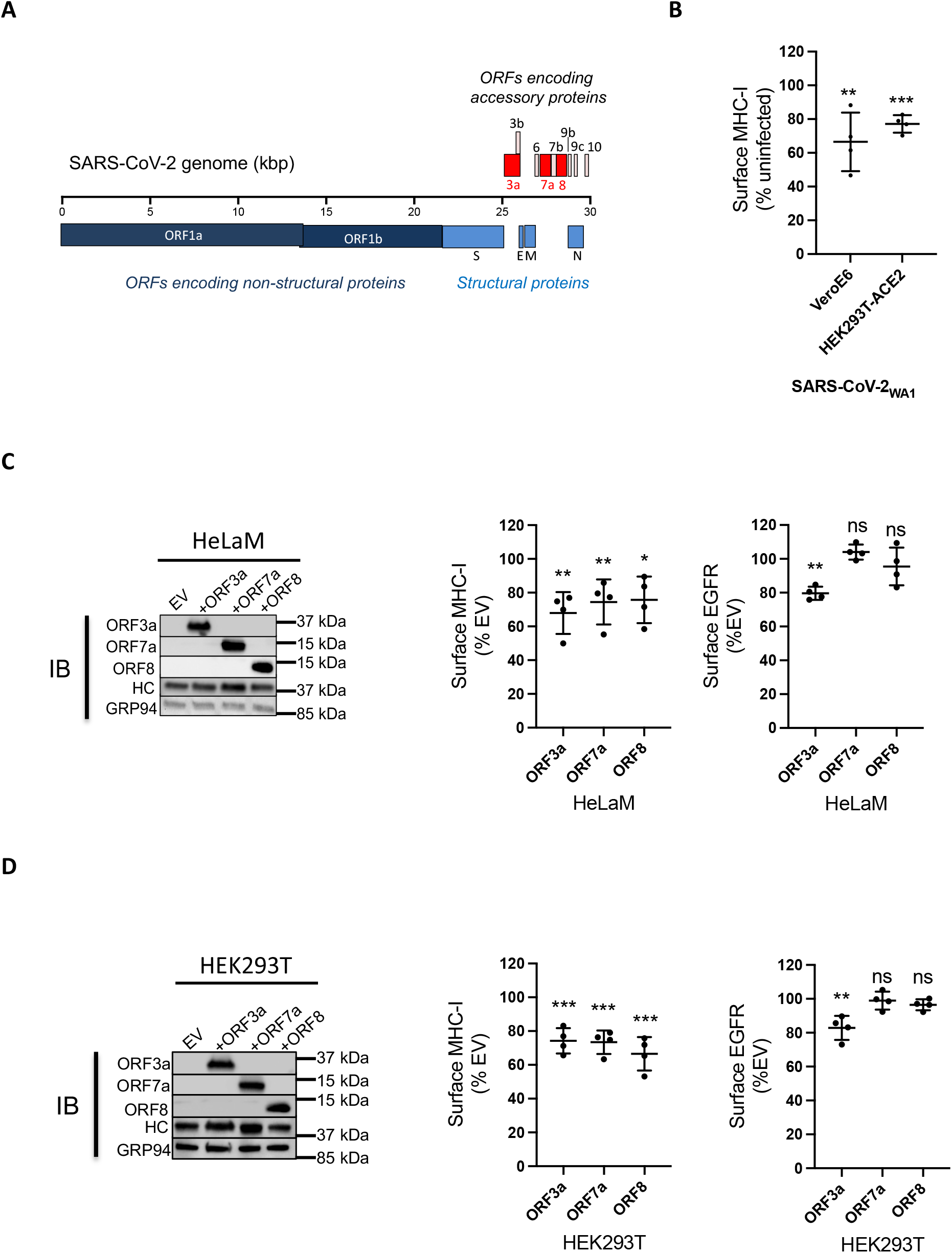
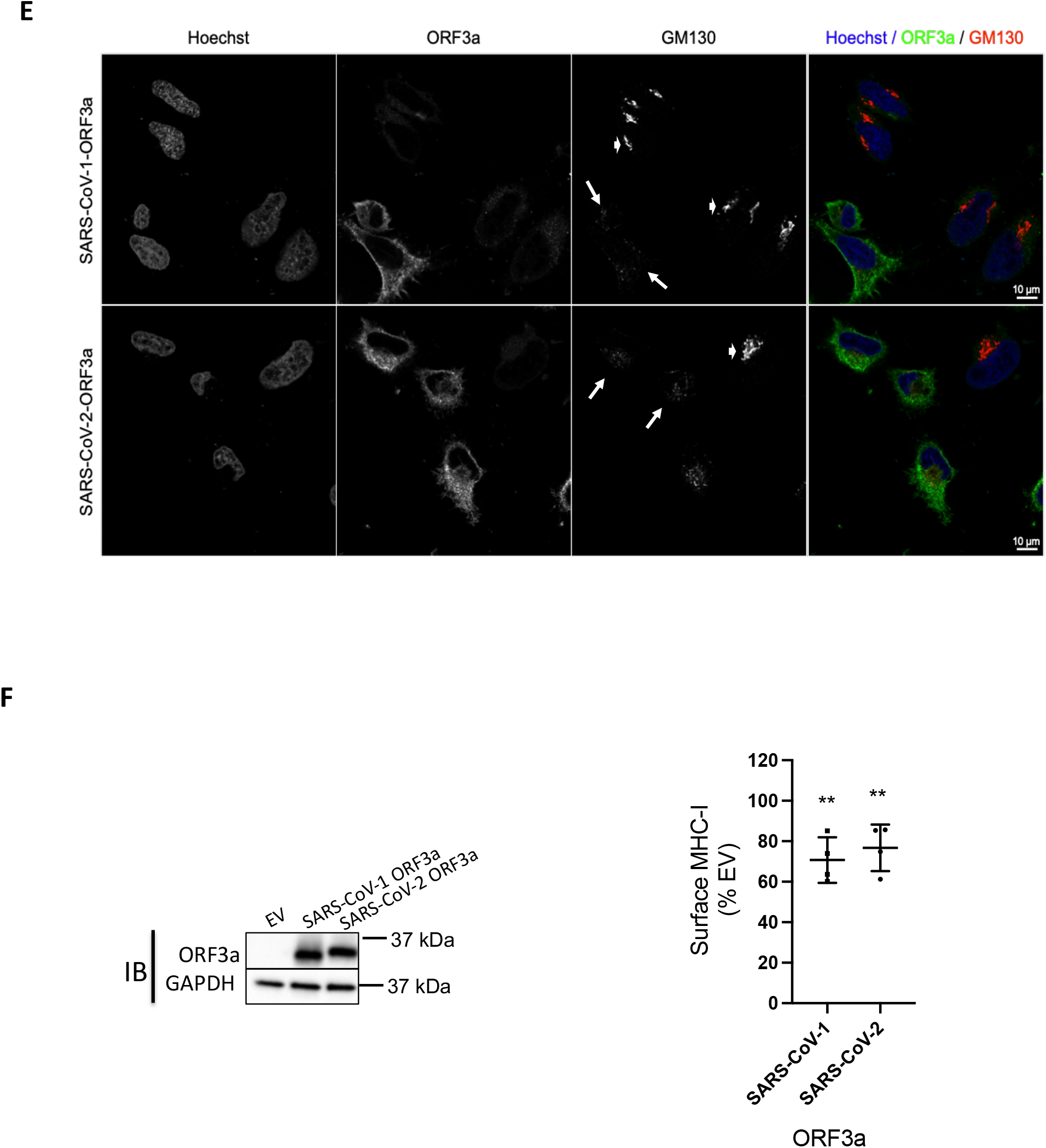
Downregulation of MHC-I by SARS-CoV-2 _WA1_ and its ER-localized gene products. (*A*) Genome organization of SARS-CoV-2. SARS-CoV-2 accessory proteins are depicted at the 3’ end of the genome. ORF3a, ORF7a and ORF8 are highlighted in red. (*B*) Vero E6 or HEK293T-Ace2 cells were infected with SARS-CoV-2_WA1_ (MOI = 10). 24 h post infection, cells were collected for flow cytometry analysis for surface MHC-I (n = 4). Mean fluorescence intensity of MHC-I staining with W6/32 was normalized to uninfected group. (*C*) Transient expression of ORF3a, ORF7a and ORF8 in HeLaM cells. Western Blot analysis of expression of accessory proteins, MHC-I heavy chain (HC) or GRP94 as a loading control, 24 h post transfection (*left*). Flow cytometric analysis of surface MHC-I (*middle*) and EGFR (*right*) in cells expressing ORF3a, ORF7a and ORF8 in HeLaM cells (n = 4). (*D*) Transient expression of ORF3a, ORF7a and ORF8 in HEK293T cells. Western Blot analysis of expression of accessory proteins, MHC-I heavy chain (HC) or GRP94 as a loading control 24 h post transfection (*left*). Flow cytometric analysis of surface MHC-I (*middle*) and EGFR (*right*) in cells expressing ORF3a, ORF7a and ORF8 in HEK293T cells (n = 4). (*E*) Effect of ORF3a on Golgi morphology. HeLaM cells were transfected with plasmids encoding SARS-CoV-1 ORF3a or SARS-CoV-2 ORF3a for 24 h. Fixed cells were stained with Hoechst (blue), for ORF3a (green), and anti-GM130 to visualize the Golgi (red). (*F*) HelaM cells were transfected with plasmids encoding SARS-CoV-ORF3a or SARS-CoV-2-ORF3a for 24 h, followed by analysis of expression by western blotting (left) or flow cytometric analysis of surface MHC-I (right, n = 4). Quantitative data shown are mean + S.D. (error bars). Statistical significance was evaluated using the unpaired Student’s t test; *p < 0.05; **p < 0.01; ***p < 0.001; ****p < 0.0001; ns, not significant.

Our analysis focused specifically on SARS-CoV-2 ORF3a and ORF7a as both of these proteins are predicted to localize to the secretory pathway and could post-translationally impact antigen presentation by MHC-I. ORF3a has a topology that suggests it can integrate into lipid bilayers (14, 15), and studies in liposomes showed that it is permeable to cations such as calcium ions indicating that it is a viroporin (15). ORF7a has an N-terminal signal peptide and one putative transmembrane domain, indicating it is membrane associated(16). While roles for ORF3a and ORF7a in regulating innate, anti-viral immune responses have been demonstrated(17), no role in adaptive immune responses has been described to date. We report here that ORF3a and ORF7a both reduce MHC-I levels to an extent comparable to ORF8 but by distinct mechanisms. ORF3a disrupts protein trafficking along the secretory pathway to inhibit surface expression of MHC-I, while ORF7a specifically interacts with the MHC-I heavy chain and delays its export from the ER.

## Results

### MHC-I is downregulated by SARS-CoV-2 infection or on expression of its ER-localized accessory proteins

To investigate the effect of viral infection on antigen presentation by MHC-I, we first looked at the effect on the surface expression of MHC-I. We infected VeroE6 cells or HEK293T cells expressing human Angiotensin-Converting Enzyme 2 or Ace2, the receptor for SARS-CoV-2, (HEK293T-hAce2) with SARS-CoV-2_WA1_ at an MOI of 10 for 24 hours, followed by estimation of surface MHC-I by flow cytometry. We observed that surface MHC-I was reduced by 30-40% in infected cells compared to uninfected cells (Fig. 1*B*, Fig. S1). To specifically look at viral proteins that might regulate MHC-I by interfering with steps in the antigen presentation pathway, we focused on those that were likely to enter the ER. These include ORF7a and ORF8, which have N-terminal ER-localization signal sequences. While ORF3a does not have such a sequence, it has been shown to be a viroporin that integrates into lipid membranes(15). These three proteins and their position in the SARS-CoV-2 genome are highlighted in Fig. 1*A*. To evaluate the effect on surface MHC-I expression, we transiently expressed them individually in the human cell lines HeLaM or HEK293T, and measured surface MHC-I 24 hours post transfection. All three proteins reduced surface MHC-I by 25-30% (Fig. 1*C,D*, Fig. S2). To ask whether this downregulation was specific to MHC-I or a consequence of an overall reduction of glycoprotein trafficking we measured the expression levels of epidermal growth factor receptor (EGFR), estimated to have a similar half-life on the surface of cells to MHC-I (~ 8-18 hours (18, 19)), and CD47, estimated to have a much longer surface half-life (~ 72 hours)(19). ORF7a or ORF8 did not affect surface expression of these proteins in either cell line, while ORF3a downregulated the expression of EGFR and CD47 in both cell lines (Fig. 1*C* (right), 1*D* (right), Fig. S3). We hypothesized that expression of ORF3a generally impairs protein trafficking through the secretory pathway, while the expression of ORF7a, like ORF8(13), specifically affects MHC-I expression.

### Expression of ORF3a induces Golgi fragmentation

After synthesis in the ER, proteins destined for the cell surface usually traverse the Golgi apparatus en route to the plasma membrane. The Golgi structure has a characteristic appearance of a stack of aligned flat cisternae, and the disruption of the Golgi morphology by pathogens can lead to the dispersal or fragmentation of the stacked cisternae and is known to block anterograde protein trafficking (20, 21). In a report that has to date not been peer-reviewed, expression of the E-protein derived from infectious bronchitis virus (IBV-E protein) or the SARS-CoV-1 ORF3a protein, was shown to disrupt Golgi morphology, with a concomitant reduction of protein trafficking to the cell surface (8). We therefore looked at the effect of SARS-CoV-2 ORF3a expression on the architecture of the Golgi apparatus to ask if the same mechanism was involved. We expressed SARS-CoV-1 ORF3a or SARS-CoV-2 ORF3a proteins in HeLaM cells and assessed Golgi morphology by confocal microscopy. In both instances we found that the Golgi was fragmented in cells expressing ORF3a, observed as a loss of signal of the Golgi marker GM130 (Fig. 1*F*, panel 3, long white arrows) compared to cells that did not express ORF3a (Fig. 1*F*, panel 3, short white arrows). Transient expression of either SARS-CoV-1 or SARS-CoV-2 ORF3a in cells for 24 hours led to a ~30% decrease in surface MHC-I levels (Fig. 1*G*). This data, along with the observed decrease in surface EGFR and CD47 (Fig. 1*C* (right), 1*D* (right), Fig. S3), leads us to propose that ORF3a downregulates surface MHC-I by blocking trafficking of glycoproteins to the cell surface by disrupting Golgi architecture.

### ORF7a slows the rate of export of MHC-I from the ER

We have successfully used HeLaM cells in the past to understand the interactions between MHC-I and the PLC(22). Therefore, to evaluate the mechanism of ORF7a-mediated MHC-I downregulation, we generated a HeLaM cell line that allowed the inducible expression of ORF7a (HeLaM-iORF7a) on treatment with doxycycline (Dox). Twenty four hours after ORF7a induction we observed a 20% decrease in surface MHC-I expression (Fig. 2*A*, left, middle). There was no change in MHC-I mRNA levels, confirming that the downregulation was not transcriptional (Fig. 2*A*, right). Since there was no overall reduction in MHC-I protein levels in HeLaM cells expressing ORF7a compared to control cells in cell lysates (Fig. 1*C*, Fig. 2*A*), we asked if the intracellular distribution of MHC-I, specifically if its trafficking through the secretory pathway, was affected by ORF7a. We took advantage of the single N-linked glycan on the heavy chain of MHC-I to assess this. As newly synthesized N-linked glycoproteins traverse the secretory pathway, glycans are modified and rendered resistant to the enzyme Endoglycosidase H (EndoH) (23). We therefore carried out a pulse-chase experiment, followed by immunoprecipitation of MHC-I and EndoH treatment to assess progress through the secretory pathway.

**Figure 2.**
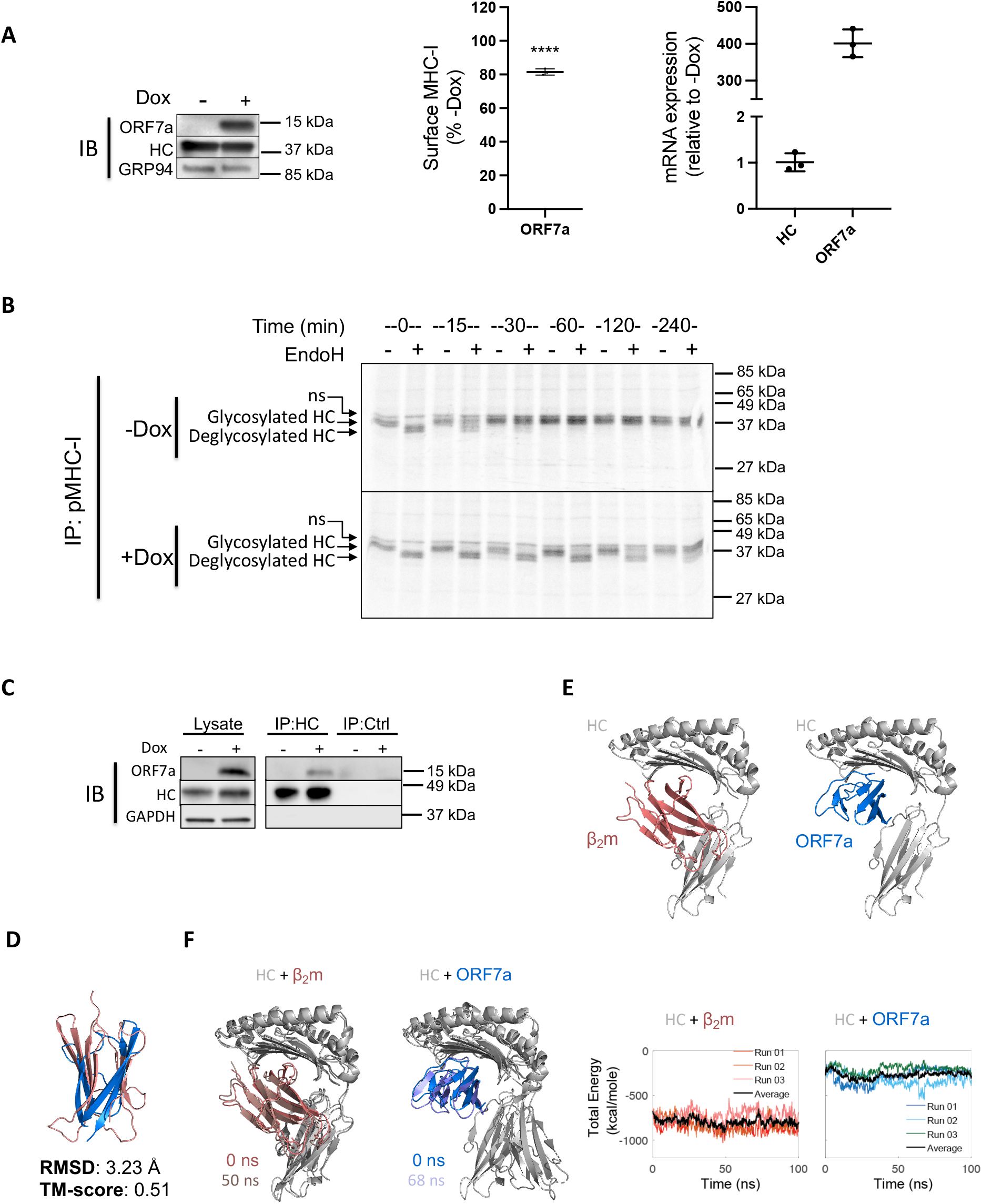
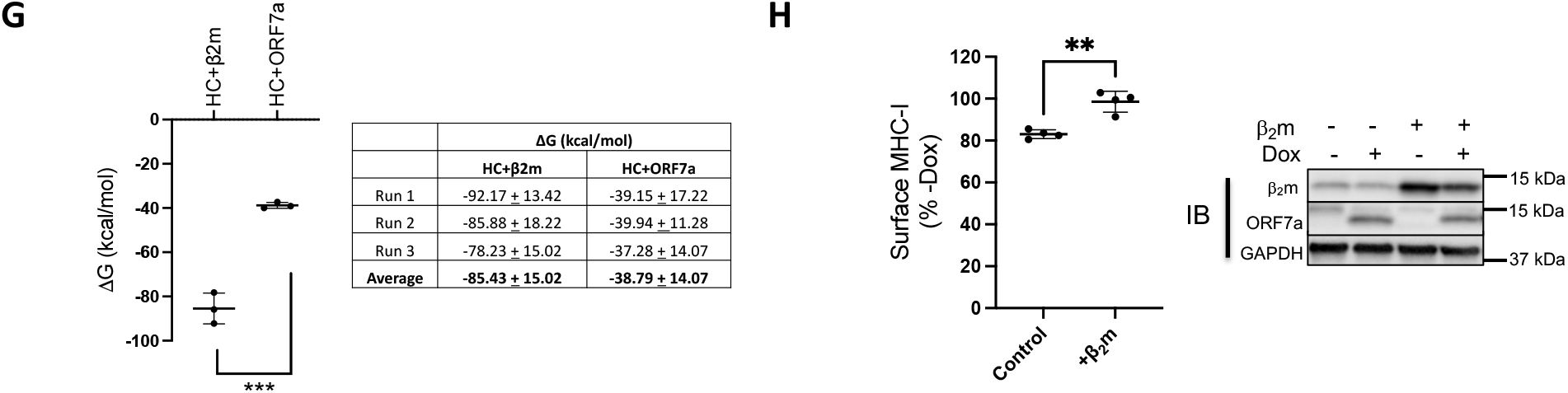
ORF7a slows the export of MHC-I from the ER by interacting with the heavy chain of MHC-I. (*A*) Stable expression of a doxycycline (Dox)-inducible construct of ORF7a in HeLaM cells was verified in uninduced and induced (+Dox for 24 h) cells by western blotting (left), and the effect on surface MHC-I was assayed by flow cytometry (middle, n = 3). MHC-I and ORF7a mRNA levels were assessed by qPCR and relative expression was normalized to uninduced (-Dox) cells (n = 3). (*B*) Uninduced (-Dox, upper panel) or induced (+Dox for 24 h, lower panel) cells were labeled with [^35^S]Met for 15 min followed by immunoprecipitation of MHC-I complexes from the lysates at the indicated time points of chase, followed by treatment with Endoglycosidase H (EndoH). Samples were visualized after separation on nonreducing SDS-polyacrylamide gels by autoradiography. (*C*) Interaction of ORF7a and MHC-I was carried out by immunoprecipitation of total heavy chain from uninduced or induced (+Dox for 24 h) cells lysed in digitonin using normal IgG as a control (IP:Ctrl) or α-HLA-A/B/C (IP: HC), followed by western blot analysis to detect ORF7a, MHC-I heavy chain (HC) or GAPDH as loading control. (*D*) Structural analysis of ORF7a and MHC-I. Alignment of the crystal structures human β_2_m in red (PDBID: 2D4F) and SARS-CoV-2 ORF7a in blue (PDBID:6W37) the RMSD (root mean square deviation) and TM-Score (template modeling score) are also reported. (*E*) Cartoon representations of the crystal structure of HLA-A2 (grey) in complex with β_2_m (red) (left panel) and the predicted structure of HLA-A2 (grey) complexed with ORF7a (blue) as modeled by ClusPro. (*F*) Cartoon representations showing a comparison of the original (0 ns) and a low energy conformer obtained from MD simulation runs (run 01, 50 ns for HC+β_2_m and run 02, 68 ns HC+ORF7a). Graphs represent total energies (van der Waal + electrostatic) of the HC+β_2_m (red, left) and HC+ORF7a (blue, right) complexes over the course of three independent MD simulation runs. Average energies are shown in black in each case. (*G*) Plot of the free energy changes of binding (ΔG) of the HC+β_2_m and HC+ORF7a complexes. (*H*) HeLaM-iORF7a cells were transiently transfected with control plasmid or plasmid encoding human β_2_m for 8 h, followed by the induction of ORF7a expression (+Dox for 24h) and effect on surface MHC-I was assayed by flow cytometry (left, n=3). Expression of ORF7a and β_2_m was assessed by western blotting, with GAPDH as loading control (right). Quantitative data shown are mean + S.D. (error bars). Statistical significance was evaluated using the unpaired Student’s t test; *p < 0.05; **p < 0.01; ***p < 0.001; ns, not significant.

Doxycycline treated and untreated HeLaM-iORF7a cells were pulse-labeled with ^35^S-methionine followed by a chase for up to 4 hours. We immunoprecipitated the MHC-I complex at different time points and incubated it with or without EndoH followed by SDS-PAGE and autoradiography. MHC-I retained in the ER is sensitive to EndoH action and treatment results in the generation of a lower molecular weight, deglycosylated heavy chain. We found that in the absence of ORF7a (-Dox), the glycan was initially sensitive to EndoH action, but acquired resistance within 30 min (Fig. 2*B*, upper panel). However, in the presence of ORF7a (+Dox), EndoH resistance was delayed by up to 4 hours (Fig. 2*B*, lower panel). This delay was specific to MHC-I as the EndoH-sensitivity pattern of the total radiolabeled glycoprotein pool, bound by the lectin Concanavalin A, was relatively unchanged by ORF7a induction after 4 hours in a similar experiment (Fig. S4).

### ORF7a competes with the β_2_m interaction with the MHC-I heavy chain

We then asked whether ORF7a was interacting with MHC-I or any components of the peptide loading machinery that might interfere with the antigen processing pathway and cause a delay in ER export. We found ORF7a co-immunoprecipitated with MHC-I heavy chain (Fig. 2*C*) but not assembled peptide-MHC-I complexes, β_2_m, or the PLC (inferred from a lack of association with tapasin). Immunoprecipitation of ORF7a to detect associated heavy chain was unsuccessful, perhaps because the ORF7a epitope recognized by the monoclonal antibody used is blocked by heavy chain interaction. These results indicate that the heavy chain can associate with either ORF7a or β_2_m but not both, suggesting that ORF7a expression may interfere with β_2_m association. We therefore analyzed the available structures of ORF7a, β_2_m, and the peptide-MHC-I complex to look for a potential interaction of ORF7a with the heavy chain. Both β_2_m and ORF7a contain an immunoglobulin (Ig)-like fold so we asked if the two are indeed structurally similar. A structural alignment using the Pairwise Structure Alignment tool provided by RCSB PDB showed a weak structural similarity, inferred by a higher root mean squared deviation (RMSD) value of 3.23 Å and a modest template modeling score (TM-score) of 0.51(Fig. 2*D*). However, this TM-score also indicates that they do have the same protein fold. We then wanted to compare the structure of the heavy chain-β_2_m complex (HC-β_2_m) with that of a likely heavy chain-ORF7a complex (HC-ORF7a). While the crystal structure is available for the former (Fig. 2*E*, left), no structural information is available for the latter. We therefore modeled the interaction of ORF7a and the heavy chain of MHC-I using the ClusPro protein-protein docking server. The predicted structure indicated that ORF7a interacts with the heavy chain in the same region as β_2_m (Fig. 2*E*, right). This is compatible with our data indicating that either ORF7a or β_2_m, but not both, can interact with the heavy chain.

We then performed all atom, explicit solvent molecular dynamics (MD) simulations to determine the free energy changes (ΔG) of binding of β_2_m and ORF7a to the heavy chain using the structural models. Analysis of the MD simulation trajectories revealed a greater Ca atom RMSD and root mean squared fluctuation (RMSF) for the heavy chain when in complex with ORF7a compared to β_2_m (Fig. S5A-C). This is reflected as a greater change in the structure and higher total energy of HC-ORF7a compared to HC-β_2_m over time during the MD simulations (Fig. 2*F*). The most stable conformation of the complex observed during the simulation vs the original structure has been superimposed (Fig. 2*F*, left), which is at 50 ns for HC-β_2_m and 68 ns for HC-ORF7a. Trajectory analysis revealed a lower total energy of HC-β_2_m compared to HC-ORF7a (Fig. 2*F*, right), which is driven largely by electrostatic interactions (Fig. S5D,E). In agreement with this, calculated ΔG values for the two complexes revealed a ~2-fold difference, which was significant (Fig. 2*H*). Based on this information, we asked if the intracellular association of ORF7a and the heavy chain could be outcompeted by overexpressing β_2_m to rescue surface MHC-I. We transiently expressed a control plasmid or a plasmid encoding β_2_m in the HeLaM-iORF7a cells, followed by induction of ORF7a expression. Cells transfected with the control plasmid resulted in the expected decrease in MHC-I surface in Dox-treated compared to untreated cells, but this was eliminated upon expression of β_2_m (Fig. 2*H*). Thus, excess β_2_m can overcome the action of ORF7a.

### Localization of ORF7a to the ER determines retention of MHC-I

Our data so far indicates the effect of ORF7a on MHC-I occurs in the ER. However, previous studies have found that ORF7a is primarily localized in the Golgi (16, 17, 24), despite harboring a dibasic amino acid sequence in its cytosolic C-terminal tail, which is a likely ER-retrieval sequence(25) (Fig. 3*A*). We therefore examined the localization pattern of the untagged ORF7a construct used throughout this study. Confocal analysis of HeLaM cells transiently expressing ORF7a showed that it localized to both the ER and Golgi (Fig. 3*B*). We generated a variant, ORF7a-ARA, where the lysine residues in the potential dibasic ER-retrieval sequence in the C-terminus of ORF7a were mutated to alanine and found that it was localized primarily in the Golgi (Fig. 3*B*). In agreement with our pulse-chase analysis, we saw that MHC-I was retained in the ER in the presence of ORF7a but this was not the case in the presence of ORF7a-ARA (Fig. 3*C*). We then investigated whether ORF7a and ORF7a-ARA impact surface MHC-I levels and antigen presentation.

**Figure 3.**
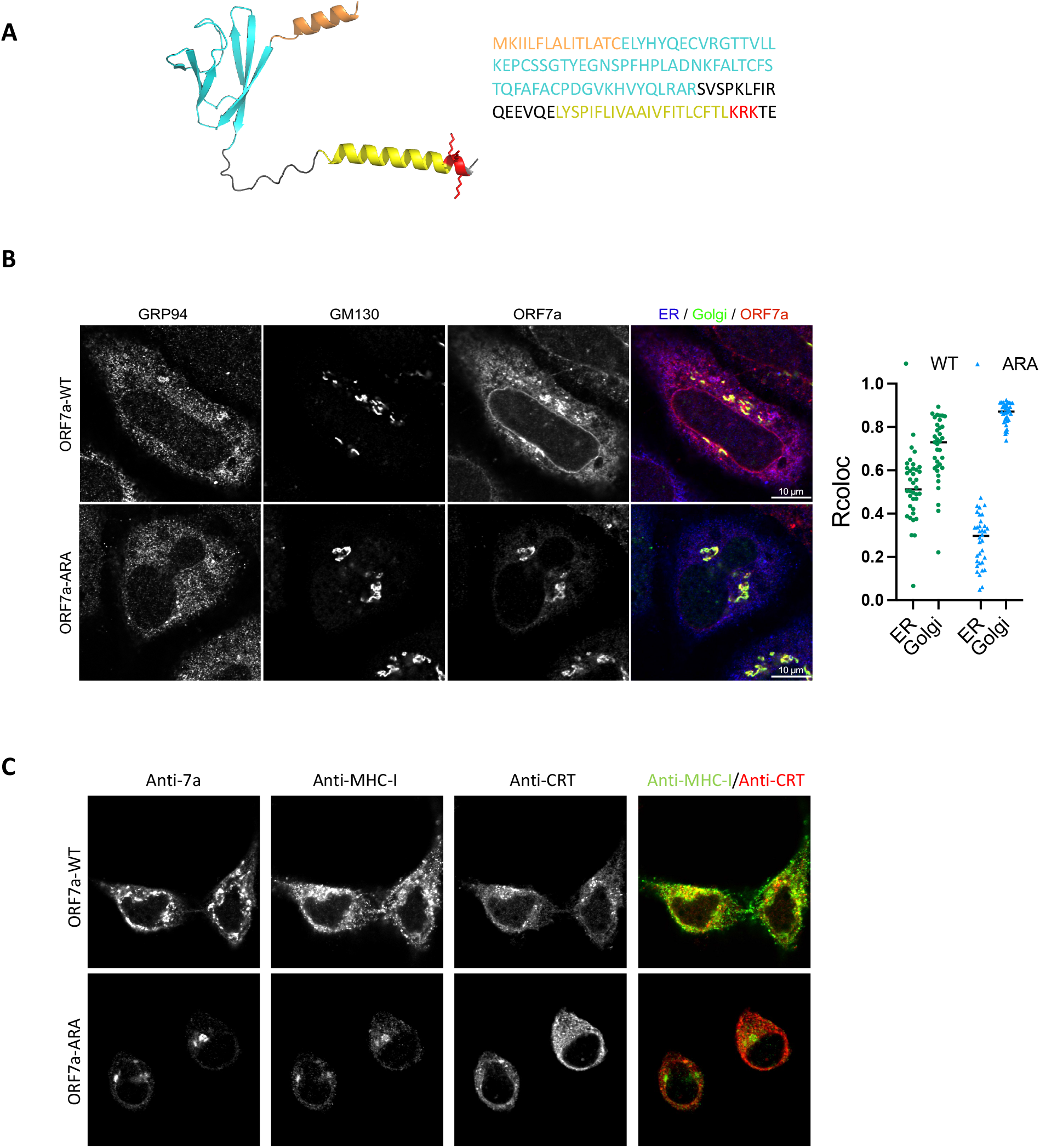
Localization of ORF7a to the ER determines retention of MHC-I. (*A*) A structural model of the full length ORF7a as generated by AlphaFold (left) and amino acid sequence of the protein (right). Color coding indicates functional features of the protein-orange, ER-targeting signal sequence; cyan, immunoglobulin-like fold; yellow, transmembrane domain; red, the ER-retrieval motif (KRK) in the cytosolic C-terminal tail with the side chains shown. (*B*) Localization of ORF7a by confocal analysis. HeLaM cells were transfected with plasmids encoding ORF7a-WT or ORF7a-ARA and 24 h post transfection cells were fixed and stained with GRP94 (ER, blue), GM130 (Golgi, green), and ORF7a, red (scale bars, 10 μm) to determine colocalization (n= 35). (*C*) Localization of MHC-I in the presence of ORF7a-WT or ORF7a-ARA was determined by transfecting HeLaM cells with plasmids encoding ORF7a-WT or ORF7a-ARA for 24 h followed by confocal analysis of ORF7a, MHC-I (green) and calreticulin (ER, red). Scale bars, 10 μm.

### ORF7a-mediated downregulation of antigen presentation by MHC-I is dependent on the ER retrieval signal

We generated HEK293T-hAce2 cells that expressed Dox-inducible versions of ORF7a or ORF7a-ARA (with the mutated ER-retrieval sequence) to assess the effect on surface MHC-I as well as antigen presentation by HLA-A*02:01 (HLA-A2), an allele that is not expressed in HeLaM cells. Following the induction of ORF7a or ORF7a-ARA expression for 24 hours (Fig. 4*A*, left), MHC-I expression on the cell surface was determined. We observed that expression of ORF7a induced an approximately 40% reduction of surface MHC-I, which was not seen upon ORF7a-ARA induction (Fig. 4*A*, middle). There was no change in MHC-I mRNA levels as assessed by qPCR analysis (Fig. 4*A*, right). We found no differences in the expression of other surface glycoproteins such as EGFR and CD47 (Fig. S6), in keeping with our previous finding that ORF7a does not affect overall glycoprotein trafficking (Fig. 1*C* (right), 1*D* (right), Fig. S3, Fig. S4).

**Figure 4.**
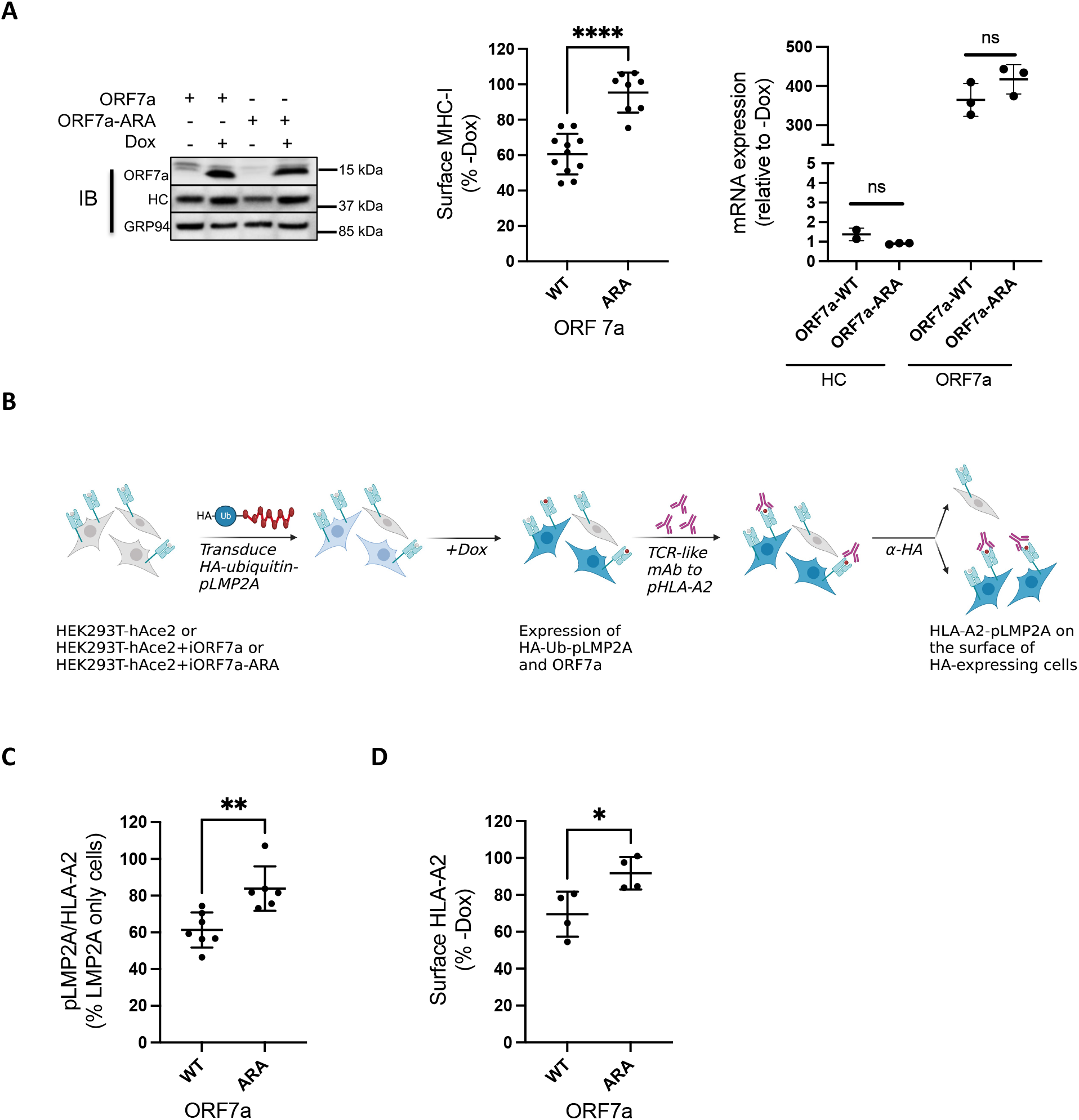
ORF7a-mediated downregulation of antigen presentation by MHC-I is dependent on the ER retrieval signal. (*A*) Stable expression of a doxycycline (Dox)-inducible construct of ORF7a-WT and ORF7a-ARA in HEK293T-hAce2 cells was verified in uninduced and induced (+Dox for 24 h) cell by western blotting (left), and the effect on surface MHC-I was assayed by flow cytometry (middle, n = 4). MHC-I and ORF7a mRNA levels were assessed by qPCR and relative expression was normalized to uninduced (-Dox) cells (n = 3). (*B*) Outline of experimental strategy employed to determine the effect of ORF7a on peptide presentation by HLA-A2. (*C*) The amount of pLMP2A presented by HLA-A2 was determined by flow cytometric analysis of induced cells (+Dox for 24 h) that expressed the fusion protein using a TCR-like monoclonal antibody specific to this complex (n = 6). (*D*) Flow cytometric analysis of surface HLA-A2 in cells expressing ORF7a-WT and ORF-7a-ARA (n = 4). Statistical significance was evaluated using the unpaired Student’s t test; *p < 0.05; **p < 0.01; ***p < 0.001; ns, not significant.

To assess whether the ORF7a-mediated decrease in MHC-I expression led to a decrease in antigen presentation, we evaluated the surface expression of a specific high-affinity peptide-MHC-I complex. The peptide we made use of is derived from the Epstein–Barr virus protein latent membrane protein 2A (LMP2A), which is presented efficiently by HLA-A2 and binds to it with high-affinity(26). We assessed the presentation of this peptide using a monoclonal antibody that mimics T cell receptors (TCRs) on the surface of CTLs that recognize this specific peptide-HLA-A2 complex(26). This flow cytometry-based assay provides a more quantitative assessment of antigen presentation than measuring cytotoxic activities of CTLs(26). We transduced HEK293T-hAce2 cells as well as these cells expressing inducible ORF7a or ORF7a-ARA (Fig. 4*B*, 4*C*) with an inducible HA-tagged ubiquitin-fusion construct encoding the LMP2A-derived peptide (pLMP2A: CLGGLLTMV). The cytosolic ubiquitin-peptide fusion protein is degraded by cellular ubiquitin hydrolases, ensuring efficient presentation of the peptide(27), and the HA-tag allows the detection of cells expressing the construct (Fig. 4*B*, Fig. S7). On induction with Dox, both ORF7a and pLMP2A were expressed, mimicking viral infection of cells with pLMP2A serving as a proxy for newly generated viral peptides. We found that in the HA-positive cells, expression of ORF7a but not ORF7a-ARA significantly reduced the presentation of pLMP2A (Fig. 4*C*). This decrease is likely due to the reduction of surface HLA-A2 mediated by ORF7a (Fig. 4*D*).

## Discussion

The SARS-CoV-2 virus is a successful pathogen that can evade both innate and adaptive immunity. Adaptive immunity includes activation of CTLs in response to viral peptides presented by MHC-I on the surface of infected cells, leading to the elimination of these cells and restriction of viral replication. Here we report two SARS-CoV-2 accessory proteins that can downregulate MHC-I levels and demonstrate that they do so by two distinct mechanisms. ORF3a is a viroporin with three transmembrane domains that can oligomerize to form an ion channel, allowing cation permeability(15). This property allows SARS-CoV-1 to disrupt the architecture of the Golgi(8) (Fig. 1*F*) and Golgi fragmentation is accompanied by a reduction of protein trafficking through the secretory pathway and a decrease in the levels of surface proteins(8). We propose this is also the case for SARS-CoV-2 ORF3a, given their similar effects on the Golgi and their high degree of similarity (85%) with 73% identity.

There is no natural virus isolate that lacks ORF3a expression, but the contribution of ORF3a to the pathogenicity of SARS-CoV-2 has been evaluated in the K18 human ACE2 transgenic mouse model(28). In this mouse model of SARS-CoV-2 infection, the pathogenicity of infectious recombinant viral particles (rSARS-CoV-2) that lacked each of the accessory proteins was evaluated. Mice infected with ΔORF3a rSARS-CoV-2 showed significantly less morbidity and mortality (75% survival) and started to recover from the virus within 5 days of infection, while mice infected with the intact virus died within 6 days. Viral loads in the nasal turbines and lungs were also significantly lower. ORF3a plays many roles that enhance viral pathogenicity in addition to the MHC-I downregulation described here, including inhibition of general surface protein trafficking, induction of apoptosis(29), inhibition of autophagy(17), and attenuating innate immune responses(17, 30). Whether a specific process or a combination of these processes reduce pathogenicity by ΔORF3a rSARS-CoV-2 remains to be evaluated.

While the downregulation of MHC-I by ORF3a seems to be a consequence of a general inhibition of protein trafficking, MHC-I downregulation by ORF7a is specific. We found that ORF7a interacts with MHC-I heavy chains in the ER and slows the export of peptide-MHC-I complexes. We were unable to detect any ORF7a interacting with tapasin, under solubilization conditions that maintain tapasin association with the other components of the PLC, or with the assembled p-MHC-I complex. We speculate that newly synthesized, free heavy chains are the targets of interaction of ORF7a. The partial structure of ORF7a has been determined (PDBID: 6W37,(31)) and it is comprised of an Ig-like domain. The human protein most structurally similar to ORF7a is intercellular adhesion molecule 3 (ICAM3), a potent immune signaling molecule. It was proposed that ORF7a may thus interact with partners of ICAM3 to modulate the immune response (32). Indeed, cell-free ORF7a can associate with the surface of uninfected CD14-positive cells (monocytes), implying a role in immune modulation although its interacting partner has not been identified(31) and how ORF7a, a transmembrane protein, might exit the cell is unclear. Human β_2_m is also an Ig-like molecule, and there is some structural overlap between β_2_m and ORF7a. Molecular modeling and MD simulation analysis of the complexes indicate that the interaction between ORF7a and MHC-I heavy chain is energetically favorable (even if it is less stable than the β_2_m and heavy chain interaction) and that the projected interaction is broadly in the same region as β_2_m. Our data also indicates that ORF7a and β_2_m compete to interact with the heavy chain as overexpressed β_2_m rescues the ORF7a-mediated downregulation of MHC-I. This suggests that ORF7a is a molecular mimic of β_2_m. Viral mimics of MHC-I heavy chain have been described, which permit infected cells to evade detection by natural killer cells that target cells with low MHC-I expression as in the case of rodent herpesvirus Peru (33) or sequester viral peptides to prevent their effective presentation as in the case of human cytomegalovirus(34). However, this report is the first description of a viral protein mimicking β_2_m to the best of our knowledge. Unlike β_2_m, ORF7a is membrane anchored and further structural and biophysical characterization of the interactions between full-length ORF7a and the heavy chain will aid in teasing out the details of this mimicry. We propose that ORF7a interacts with the heavy chain to interfere with β_2_m association, which delays the formation of heavy chain-β_2_m heterodimers, impairs their recruitment to the PLC, and reduces viral peptide loading and ER export (Fig. 5*A*). We have no evidence to suggest ORF7a directly interferes with peptide loading; the reduced surface presentation of the LMP2A-derived peptide by HLA-A2 may simply be a consequence of the allele’s reduced surface expression. A comprehensive analysis of the immunopeptidome of cells expressing ORF7a vs ORF7a-ARA will properly evaluate the full impact on peptide presentation.

**Figure 5.**
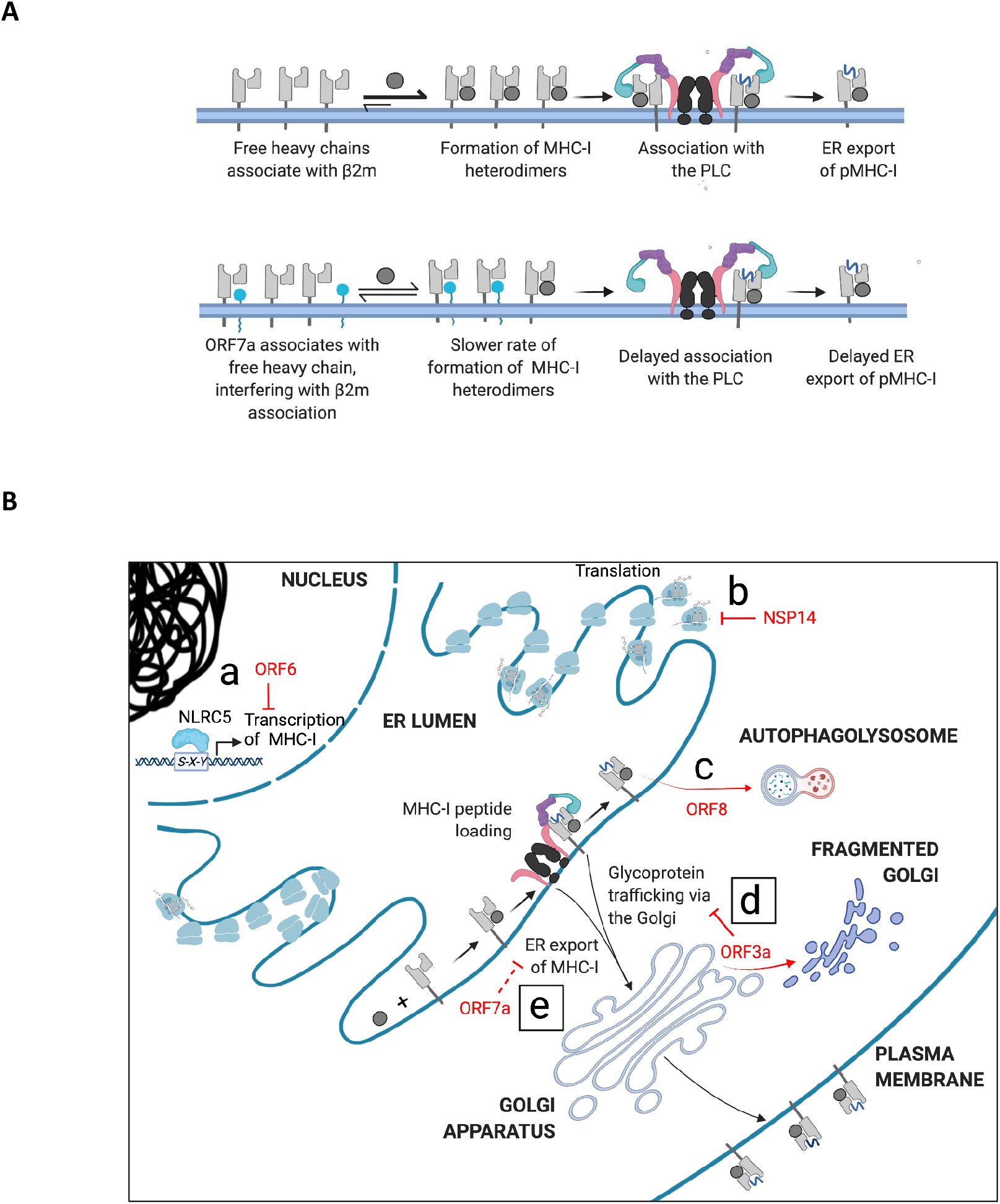
Mechanisms by which SARS-CoV-2 evades adaptive immune responses mediated by MHC-I (*A*) Proposed mechanism by which ORF7a slows the export of MHC-I from the ER. In uninfected cells the free heavy chain associates with β_2_m to form the MHC-I heterodimer. This associates with the peptide loading complex (PLC) and MHC-I molecules loaded with peptide (pMHC-I) are released for export from the ER en route to the cell surface (upper panel). ORF7a interacts with free heavy chain, interfering with its association with β_2_m and thus the peptide loading complex, delaying exit from the ER (lower panel). (*B*) Various gene products of SARS-CoV-2 downregulate MHC-I-mediated immune responses in an infected cell. ORF6 prevents the transcriptional upregulation of MHC-I following infection (a), while NSP14 globally downregulates translation (b). ORF8 interacts with HLA-A2 and targets it to the autophagosome where it is destined to be degraded on fusion with lysosomes to form the autophagolysosome (c). We report that expression of ORF3a causes Golgi fragmentation, inhibiting the trafficking of MHC-I and other surface proteins (d), and that ER-localized ORF7a impedes the export of MHC-I from the ER, downregulating surface MHC-I (e).

The ORF7a-induced downregulation of surface MHC-I and antigen presentation was alleviated when the ER-retrieval sequence of ORF7a was mutated, causing it to localize to the Golgi. In all our experiments we expressed untagged ORF7a and found that it localized to both the ER and the Golgi by confocal microscopy. This is in contrast to three previous studies that indicated ORF7a was primarily localized to the Golgi apparatus (16, 17, 24). The ORF7a constructs used in those studies were tagged C-terminally with either a small FLAG-tag (8 aa), or 2x StrepII tag (30 aa), or mCherry-FLAG (244 aa). The function of the di-basic ER retrieval signal is likely to be masked by C-terminal extensions, resulting in a cellular distribution similar to the ORF7a-ARA mutant. One study of untagged SARS-CoV-1 ORF7a indicated that it localized to the Golgi and weakly to the ER in Vero cells(35, 36), which is more in keeping with our observations. The two CoV-1 and CoV-2 proteins show a sequence identity of 87% and the ER retention sequence is conserved between them(31). Given the functional consequence of localization of ORF7a we have observed, we suggest that studies employing C-terminally tagged ORF7a need to be interpreted with caution. However, a recent report that has not been peer-reviewed found that C-terminally tagged ORF7a was capable of downregulating MHC-I in HEK293T cells(37). While we expressed ORF7a-ARA in an inducible manner, the tagged ORF7a was transiently expressed, and we are investigating whether this or other mechanisms account for this difference.

Unlike ORF3a, SARS-CoV-2 variants lacking functional ORF7a appear to exist in nature. A viral strain with a 392 nucleotide deletion resulted in the expression of the signal sequence of ORF7a in fusion with ORF8, and another strain carrying a deletion of 227 nucleotides resulted in the fusion of ORF7a and ORF7b(38). Thus both strains are effectively ORF7a knockouts, but no study to date has determined if this changes disease severity. However, a strain that carried a truncated ORF7a caused by a 115 nucleotide deletion exhibited impaired in vitro growth and led to elevated interferon-mediated responses. In addition, it died out in the immunocompetent population 1.5 months after it was detected(39). Another study identified a mutation in the transmembrane domain of ORF7a that improved its stability and caused severe infection despite low viral loads(40) although the mechanism behind this has not been determined. A mouse model of COVID19 using ΔORF7a rSARS-CoV-2 showed that this virus had a replication defect in vitro, was cleared more efficiently from the nasal turbines than intact virus, and lung lesions caused by it were also resolved sooner(28). However, although the mice began to recover earlier than the mice infected with the intact virus, only 25% survived. This is in contrast to the 75% that survived when ORF3a is deleted. As with ORF3a, it is difficult to parse out which function of ORF7a is responsible for amelioration of symptoms on infection with ΔORF7a rSARS-CoV-2. As a cautionary note, a similar analysis with ORF8 indicated that ΔORF8 rSARS-CoV-2 was as virulent as the WT virus in mice, but population studies in humans of SARS-CoV-2 with a 382 nucleotide deletion in ORF8 show that it results milder infection and is not fixed in the population(41).

We assessed the effect of ORF7a on antigen presentation by detecting the specific HLA-A2/pLMP2A complex and found that it was downregulated in the presence of ORF7a but not ORF7a-ARA. On inducing the expression of the LMP2A-derived peptide at the same time as ORF7a expression, we are mimicking the events of an infection where ORF7a is expressed along with pLMP2A, which serves as a proxy for viral peptides that may be presented by the cell. The interaction of ORF7a with free heavy chains likely delays the loading of MHC-I viral peptides by the PLC and enables the virus to evade immune detection. To evaluate the effect of ORF7a on MHC-I in the context of infection we used the recombinant SARS-CoV-2 mNeonGreen (CoV-2-mNG) virus, where ORF7a has been replaced with the fluorescent protein, rendering it a knockout for ORF7a(42). However, we found no difference in surface MHC-I levels in cells infected with this virus compared to the wildtype (Fig. S8). There are multiple factors that may explain this. First, many SARS-CoV-2 proteins contribute to the downregulation of MHC-I on infection: transcriptional suppression of MHC-I expression upon infection by accessory protein ORF6(12), inhibition of global translation by the non-structural protein NSP14(43); ORF3a and ORF8(13) also contribute to post translational downregulation of MHC-I. Second, ORF7a is expressed abundantly early in infection (within 3 hours of the 8 hour replication cycle(44)), presumably targeting nascent MHC-I molecules to prevent the presentation of peptides produced early in the viral infection, while our analysis was carried out 24 hours post infection, perhaps missing the window of ORF7a-specific effects. Finally, rather than the levels of surface MHC-I, antigen presentation needs to be assessed at the early timepoint to see if there is indeed an inhibition of viral peptide presentation, as is suggested by our data in Fig. 4*E*.

Given the success of SARS-CoV-2 as a pathogen, it is unsurprising that it has evolved multiple strategies for evading immune responses. Many studies have focused on the role of SARS-CoV-2 proteins in regulating innate immunity(17, 45) and the mechanisms of how adaptive immune responses, specifically T cell responses, are modulated by the manipulation of MHC-I, are now coming to light. There are several regulatory points in the antigen processing pathway where MHC-I expression may be altered by SARS-CoV-2. Here we describe two additional viral proteins that manipulate MHC-I levels post-translationally by distinct mechanisms. There is a global decrease in trafficking of surface proteins caused by ORF3a, and a specific retention of MHC-I in the ER by ORF7a. Both events delay MHC-I trafficking and impair antigen presentation, (Fig. 5*B*). As with many viruses, these accessory proteins are multi-functional. A deeper understanding of the interactions of SARS-CoV-2 proteins with host proteins that manipulate immune responses will help in elucidating the mechanisms that contribute to pathogenicity and provide new avenues for the development of therapeutics, as well as aid in our understanding of any related emergent pathogens in the future.

## Materials and Methods

### Plasmids

pLEX307-ACE2-blast was a gift from Alejandro Chavez & Sho Iketani (Addgene plasmid # 158449). pCAGGS vector encoding untagged viral proteins ORF3a, ORF7a, ORF8 and ORF7a-ARA were generated by Gibson assembly by PCR amplification from their respective HA-tagged constructs in pCAGGS(43). Plasmid pTRIPZ-puro-ORF7a-WT or ARA was generated by Gibson assembly by the PCR amplification of the genes of interest from their respective pcAGGS constructs and pTRIPZ-puro vector backbone digested with AgeI and EcoRI according to the manufacturer’s instructions. pTRIPZ-puro-HA-Ub was generated by Gibson assembly by PCR amplification of HA-tagged ubiquitin gene from the plasmid HA-Ubiquitin which was a gift from Edward Yeh (Addgene plasmid # 18712) and pTRIPZ-puro digested with AgeI and XhoI. This plasmid was digested with EcoRI, and was used along with single stranded DNA primers encoding pLMP2A with 20 bp overlap with the vector in a Gibson assembly reaction according to the manufacturer’s instructions to generate pTRIPZ-puro-HA-Ub-pLMP2A. All constructs were verified by sequencing. The details of primers used are listed in Table S1.

### Antibodies

Rat anti-human GRP94 (ADI-SPA-850) was purchased from Enzo Life Sciences. The antihuman tapasin antibodies PaSta1(46) and the anti-HLA-A2 antibody BB7.2(47) have been reported previously. We used mouse anti-human HLA-ABC (W6/32) conjugated to Alexa Fluor® 647 (MCA81A647, Bio-Rad), which is known to also react with MHC-I of African Green monkey from which Vero E6 cells are derived(48) for flow cytometric analysis. Mouse anti-human EGFR antibody conjugated to Alexa Fluor® 647(352917), mouse anti-human CD47 antibody conjugated to APC (323123) and anti-HA Alexa Fluor® 488 anti-HA.11 Epitope Tag (901509) were purchased from Biolegend. The antibody to ORF3a was a gift from Prof. Carolyn E. Machamer(8), and antibodies to ORF7a (GTX632602) and ORF8 (GTX135591) were purchased from GeneTex. We used a mouse monoclonal antibody to GM130 (BDB610822, from BD Sciences) and a rabbit polyclonal antibody to GM130 (11308 from Proteintech) for Golgi staining. The HLA-ABC antibody (clone YTH862.2) was purchased from Novus for immunoprecipitation and immunofluorescence imaging of MHC-I heavy chains. The fluorescently labeled spike protein antibody was purchased from eBioscience (53-6491-82). The TCR-like mAb that detects HLA-A2-pLMP2A (CLGGLLTMV) complexes by flow cytometry was a gift from Prof. Paul A. MacAry (41). All secondary antibodies used for flow cytometry and immunofluorescence imaging were purchased from Invitrogen, and those for immunoblotting that were coupled to horseradish peroxidase were purchased from Jackson ImmunoResearch.

### Reagents

Transit®-2020 and Transit®-Lenti (Mirus), Polybrene (Millipore), Triton X-100 (AmericanBio), Digitonin (Calbiochem), EDTA-free protease inhibitor mixture (Roche Applied Science), L-[^35^S]methionine, and L-[^35^S]cysteine (PerkinElmer Life Sciences), protein molecular weight markers, [Methyl-^14^C] Methylated (PerkinElmer Life Sciences), TRIzol (Invitrogen), ConcanavalinA-Sepharose 4B, Restore™ PLUS Western blotting stripping buffer (Thermo Scientific) and doxycycline hyclate (Sigma).

### Transfection and transduction

For DNA plasmid transfection, cells were plated one day before and transfected using Transit®-2020 according to the manufacturer’s instructions. Lentiviral particles were generated in HEK293T cells transfected with the packaging vector pCL-Ampho (Imgenex) and the construct of interest using Transit®-Lenti (Mirus) in Opti-MEM (Invitrogen), followed by two rounds of transduction of the recipient cell line with 8 μg/ml Polybrene (Millipore). Cells stably expressing desired construct were obtained by selection in complete medium supplemented with 2 μg/ml puromycin (Sigma) or 10 μg/ml blasticidin.

### Cell culture

VeroE6, HeLaM and HEK293T cells and derivatives were grown in Dulbecco’s modified Eagle’s medium (Sigma) supplemented with 10% fetal calf serum (HyClone) and 1% penicillin/streptomycin (Invitrogen).

### Cell lines generated

HEK293T cells expressing human Ace2 (HEK293T-hAce2) were generated by lentiviral transduction of cells with pLEX307-ACE2-blast, the vector encoding human ACE2, followed by selection with balsticidin (10 ug/ml). Expression of hAce2 was confirmed by western blotting. Inducible cell lines: HeLaM cells expressing inducible ORF7a (HeLaM-iORF7a) or HEK293T-hAce2 cells expressing inducible proteins (HEK293T-hAce2+iORF7a or HEK293T-hAce2+iORF7a-ARA) were generated by lentiviral transduction of the plasmid encoding the protein of interest followed by puromycin selection (2 ug/ml). HEK293T-hAce2 cell line and its variants were again transduced with a plasmid encoding the fusion protein of HA-tagged ubiquitin and LMP2A derived peptide (HA-Ub-pLMP2A) to generate HEK293T-hAce2+ipLMP2A, HEK293T-hAce2+iORF7a+ipLMP2A, HEK293T-hAce2+iORF7a-ARA+ipLMP2A. Since this construct also carried the puromycin marker, transduced cells expressing the HA-Ub-pLMP2A construct were identified by intracellular staining with anti-HA as outlined in Fig. 4*B*. In all instances, induction of protein expression was carried out using 5 μg/ml doxycycline for 24 hours.

### SARS-Cov-2 Infection

High-titer stocks of SARS-CoV-2 virus (isolate USA-WA1/2020, CoV-2 WA1) obtained from BEI reagent repository and SARS-CoV-2-mNG(42) were obtained by passage in Vero E6 cells. Viral titers were determined by plaque assay on Vero E6 cells. Virus was cultured exclusively in a biosafety level 3 facility. Vero E6 cells were either mock infected or infected at a multiplicity of infection (MOI) of 10 in serum free DMEM for 1 h. Unbound virus was washed off, and cells were maintained in 2% FBS/DMEM at 37°C for 24 hours, followed by staining for surface MHC-I for flow cytometric analysis. Poststaining cells were fixed using Cytofix Cytoperm and intracellular FACS was carried out where indicated.

### Flow Cytometry

Cells were detached with trypsin, and sequentially stained with primary and secondary antibody (unless the primary antibody was fluorescently labeled) in FACS staining buffer (PBS + 0.5% BSA + 0.02% sodium azide) for 1 h at 4 °C. Cells were washed three times with FACS buffer after incubation with primary and secondary antibody, and analyzed using FACSCalibur (BD Biosciences) and FlowJo software (Tree Star). Staining with propidium iodide was carried out to distinguish dead from live cells in unfixed cells, and the Live/Dead Fixable Red Dead Cell kit (L34971, Invitrogen) was used in experiments that required fixing and intracellular staining. To distinguish transfected vs untransfected cells, intracellular staining for the protein of interest was carried out after surface staining for MHC-I. Cells were fixed with 4% PFA (or CytofixCytoperm for virus infected cells), followed by cell permeabilization with 0.4% TritonX-100 in in intracellular FACS buffer (TBS + 2.5% BSA +0.1% Tween20), followed by staining with antibodies in intracellular FACS buffer overnight at 4 °C. Cells were then washed with TBS followed by staining with secondary antibodies or analysis as mentioned above.

### Co-immunoprecipitation

HeLaM cells were lysed in TBS, pH7.4, containing1% digitonin + protease inhibitors + 2 mM CaCl2 at 4 °C for 1 h. Clarified lysates were then either prepared for SDS-PAGE by the addition of Laemmli sample buffer or immunoprecipitated with indicated antibodies plus protein G-Sepharose for 2 h at 4 °C. After three washes in 0.1% digitonin lysis buffer and elution in nonreducing sample buffer, immunoprecipitates were separated by SDS-PAGE under non-reducing conditions followed by western blotting.

### Immunoblotting

Cell lysates were prepared by directly lysing an equal number of cells in 2% Laemmli sample buffer. Proteins were separated by SDS-PAGE followed by Western blot analysis. After probing with primary antibodies in TBST (TBS+ 0.1% Tween20) + 2.5% BSA overnight at 4 °C, blots were incubated with secondary horseradish peroxidase-conjugated antibodies in TBST + 0.5% BSA for 1 h at RT, and analyzed by the Chemidoc MP Imaging System (Bio-Rad). Blots were stripped and re-probed following treatment with Restore™ PLUS Western blotting stripping buffer according to the manufacturer’s instructions. Data are representative of experiments carried out with at least three biological replicates.

### Immunofluorescence Assay

Visualization of proteins of interest was carried out by confocal microscopy. HeLaM cells were plated on coverslips 24 hours before transfection with plasmids encoding DNA of interest and processed 24 hours post transfection. Cells were washed with PBS and fixed with 4% PFA for 15 min at room temperature, rinsed thrice with TBS + 100 mM glycine, permeabilized with 0.4% TritonX-100 in TBST + 2.5% BSA for 10 min. Antibody staining was carried out in intracellular FACS buffer overnight at 4 C in a humidified chamber, followed by 3 washes with TBST and incubation with secondary antibodies conjugated to fluorescent dyes and Hoechst 33342 stain for 30 min at 4 C and 3 more washes with TBST. Coverslips were then mounted on slides using ProLongGold mountant according to the manufacturer’s instructions followed by imaging on the Leica stellaris 8. Images are representative of at least 3 biological replicates and at least 10 fields of view for each. Any post-acquisition processing was kept consistent across images. The images were analyzed by Fiji(49). By applying the Polygon Selection function to GRP94 stained images, the cells were segmented to define the region of interest. Colocalization of ORF7a with GRP94 and GM130, respectively, were analyzed by Colocalization Threshold function of Fiji. The Rloc values which represent the Pearson’s coefficient above the threshold were plotted.

### Quantitative RT-PCR

Cells were harvested in TRIzol, and RNA was extracted using the Direct-zol RNA MiniPrep kit (Zymo Research). cDNA synthesis was carried out using the iScript™ cDNA synthesis kit (Bio-Rad). PerfeCTa® SYBR® Green SuperMixMaster Mix (QuantaBio) was used for qPCR analysis according to the manufactures’ instructions, and the reactions were set up by the epMotion M5073 robot (Eppendorf) and carried out in the CFX384 Touch™ real-time PCR detection system (BioRad). The details of primers used are listed in Table S2.

### Metabolic labeling and immunoprecipitation

HeLaM-iORF7a cells were incubated in cysteine- and methionine-free Dulbecco’s modified Eagle’s medium supplemented with 5% dialyzed fetal bovine serum (Invitrogen), 1% Glutamax, 1% penicillin/streptomycin for 1 h, followed by labeling with [^35^S]methionine/cysteine (PerkinElmer Life Sciences) in the same medium for 15 min. Cells were then spun down and resus-pended in complete medium, and cells were collected at indicated times of chase. Cell lysates were prepared in 1% Triton X-100 in TBS (10 mM Tris, pH 7.4, 150 mM sodium chloride) + protease inhibitors and precleared with normal rabbit IgG, followed by immunoprecipitation with W6/32 and protein G-Sepharose. The immunoprecipitates were washed three times with TBS + 0.5% Triton X-100, and proteins were visualized by autoradiography after separation on nonreducing SDS-polyacrylamide gels.

### Structural analysis

The crystal structures deposited in the protein data bank (PDB) of human β_2_m (PDBID:2D4F)(50), ORF7a (PDBID:6W37)(51), HLA-A2/β_2_m/peptide complex (PDBID: 1DUY)(52) were used in the analysis. The structural alignment of hβ_2_m and ORF7a was carried out using the Pairwise Structure Alignment tool provided by RCSB PDB(53, 54), using the TM-align parameter. The molecular docking of ORF7a and HLA-A2 heavy chain was carried out using ClusPro(55–58) and analysis of the interactions was carried out using the PRODIGY server (59, 60). The model of the full length of ORF7a was generated using AlphaFold(61).

### Molecular dynamics simulations and binding free energy analysis

Nanoscale Molecular Dynamics (NAMD) V2.13 software (62) and CHARMM36 force field (63) were utilized to perform the MD simulations as described previously (64–66). The simulation systems consisting of the biomolecular complexes of HC+β_2_m and HC+ORF7a described above were generated using the QwikMD Toolkit (67) available as a plugin in Visual Molecular Dynamics (VMD) software V1.9.3 (68). Briefly, TIP3P (69) cubic water box was used to explicitly solvate the proteins. Sodium chloride (NaCl) at concentration of 0.15 M in the explicit solvent was used for charge neutralization with Periodic Boundary Conditions applied. Prior to performing MD production runs, energy minimization was carried out on the biomolecular simulation systems for 2000 timesteps (0.004 ns) followed by an annealing step for 0.25 ns where temperature was elevated at 1 K increments from 60 K to 310 K. Temperature and pressure were then held constant at 310 K using Langevin temperature control and at 1.0 atm using Nose-Hoover Langevin piston control, respectively, followed by 1 ns equilibration step where backbone atoms of proteins were constrained through harmonic potential. Finally, three independent 100 ns runs were performed for each of the HC+β_2_m and HC+ORF7a complex biomolecular simulation systems. A 2 fs integration time step was selected for all simulations where trajectory frames were saved every 10000 steps. A 12 Å cut-off with 10 Å switching distance was chosen to handle short-range non-bonded interactions, while long-range non-bonded electrostatic interactions were handled using Particle-mesh scheme at 1 Å PME grid spacing. Analysis of interaction energy (vdW and electrostatic) throughout the 100 ns simulation was performed using “NAMD Energy” analysis tool available as part of VMD (68). Binding free energy changes (ΔG) were estimated through molecular mechanics Poisson-Boltzmann surface area (MM-PBSA) method (70) using CaFE 1.0 tool (71) combined with VMD (68) over the course of entire 100 ns MD simulation runs. All structures were visualized in PyMOL and images were created in PyMOL (The PyMOL Molecular Graphics System, Version 2.0 Schrödinger, LLC).

## Acknowledgments and funding sources

We thank Dr. Carolyn Machamer of Johns Hopkins University School of Medicine for advice on the function of ORF3a and for a critical reading of the manuscript. M. L-R. was supported by grant number KL2 TR001862, J.P. was supported by 5-T32 HL 7975-19, and J. C-C. was supported by a Cancer Research-Irvington Postdoctoral Fellowship. The work was supported by NIH grant RO1 AI059167 to P.C. W.S.A. is supported by a scholarship from the College of Health & Life Sciences, Hamad Bin Khalifa University, a member of the Qatar Foundation. Some of the computational research work reported in the manuscript were performed using high-performance computer resources and services provided by the Research Computing group in Texas A&M University at Qatar. Research Computing is funded by Qatar Foundation for Education, Science and Community Development (http://www.qf.org.qa).

