## Supplementary Information for "SARS-CoV-2 accessory proteins ORF7a and ORF3a use distinct mechanisms to downregulate MHC-I surface expression"

**This PDF file includes:**

Figures S1 to S8
Tables S1 to S2

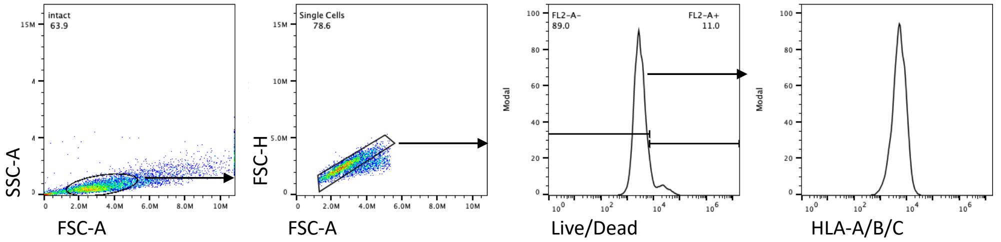


**Fig. S1:** Representative gating strategy for FACS analysis of MHC-I (HLA-A/B/C) in SARS-CoV-2$_{WA1}$ -infected Vero E6 or HEK293T-hACE2 cells.

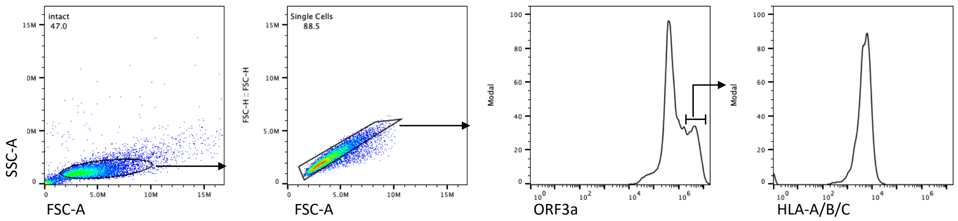


**Fig. S2:** Representative gating strategy for FACS analysis of MHC-I (HLA-A/B/C) in cells transfected with accessory proteins, with ORF3a as an example.

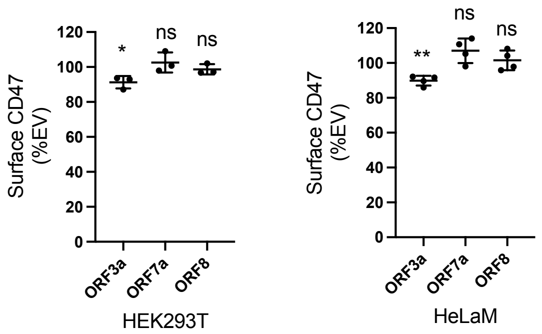


**Fig. S3:** Expression of surface glycoprotein CD47 is downregulated by ORF3a. Transient expression of ORF3a, ORF7a and ORF8 in HEK293T (n = 3) or HeLaM (n = 4) cells was carried out followed by flow cytometric analysis of CD47 expression 24 h post transfection. Quantitative data shown are mean $\pm$ S.D. (error bars). Statistical significance was evaluated using the unpaired Student's t test; *$p < 0.05$; **$p < 0.01$; ***$p < 0.001$; ****$p < 0.0001$; ns, not significant.

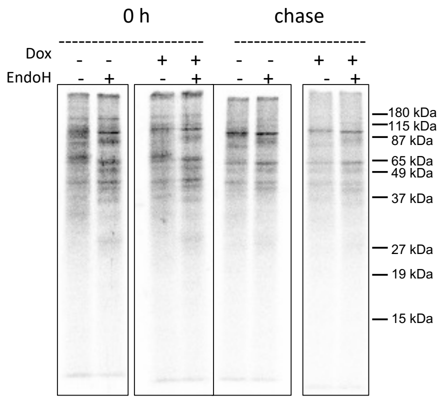


**Fig. S4:** Overall glycoprotein export is not affected by the expression of ORF7a. Uninduced (-Dox) or induced (+Dox for 24 h) cells were labeled with [$^{35}$S]Met for 15 min followed by the pull down of glycoproteins using the lectin Concanavalin A (ConA) conjugated to beads from the lysates at 0 h and 4 h of chase, followed by treatment with endoglycosidase H (EndoH). Samples were visualized after separation on nonreducing SDS-polyacrylamide gels by autoradiography.

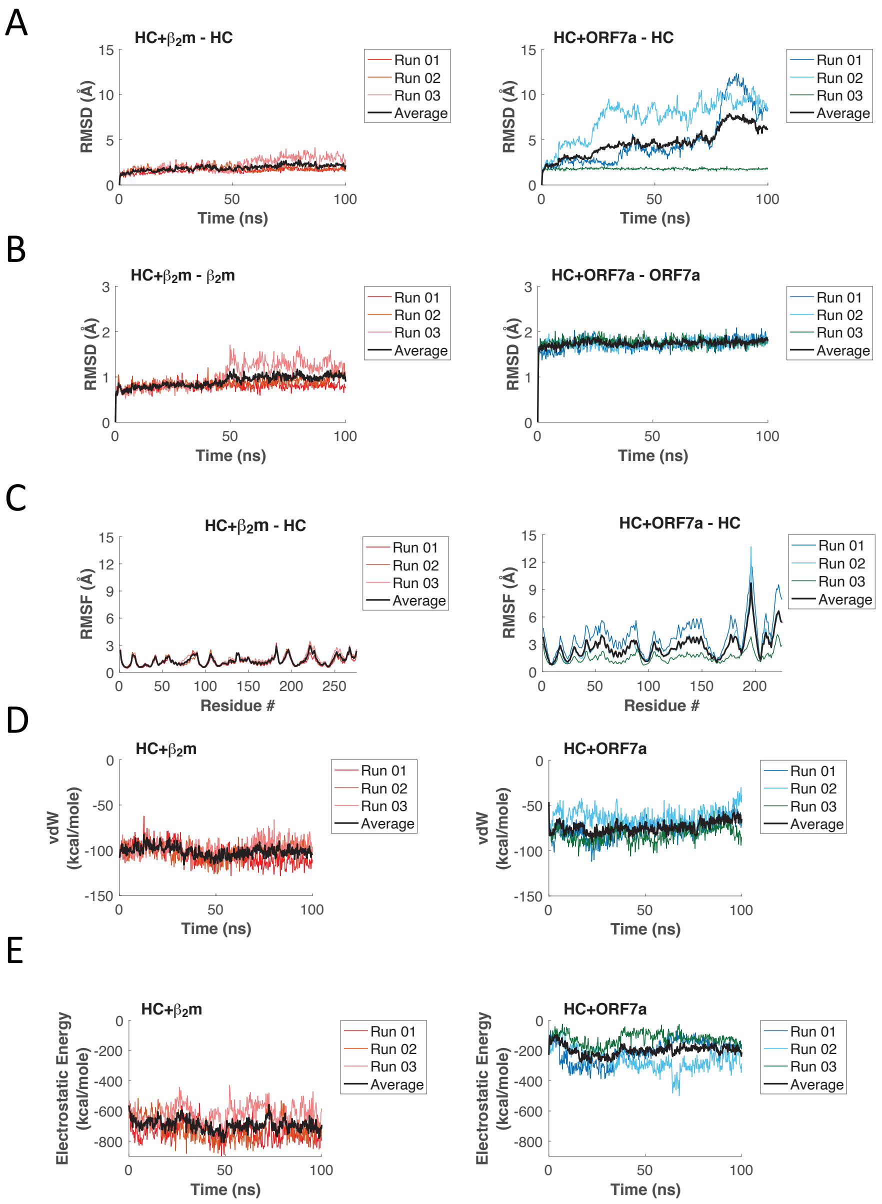


**Fig. S5:** Analysis of MD simulation trajectories. (A,B) Graphs showing root mean squared deviation (RMSD) of Cα atoms of the heavy chain (HC) (A) and either β2m (left panel) or ORF7a (right panel) (B) in the HC+β2m and HC+ORF7a complexes over three independent 100 ns simulations. (C) Graphs showing root mean squared fluctuation (RMSF) of Cα atoms of HC when associated with either β2m (left panel) or ORF7a (right panel) over three independent 100 ns simulations. (D,E) Graphs showing van der Waal (D) and electrostatic (E) energies of the HC+β2m (left panel) and HC+ORF7a (right panel) over the course three independent 100 ns simulations. Black lines in each graph show the average values of the indicated parameter calculated from all three MD simulation runs.

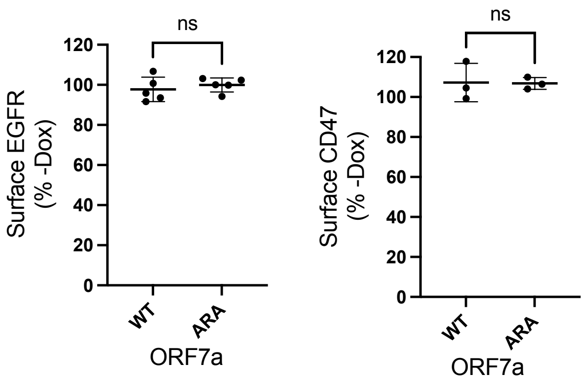


**Fig. S6**: Flow cytometric analysis of surface EGFR (left) and CD47 (right) levels in induced HEK293T-hAce2 cells+ORF7a-WT and HEK293T-hAce2 cells+ORF-7a-ARA cells (+Dox for 24 h), plotted relative to uninduced cells (-Dox).

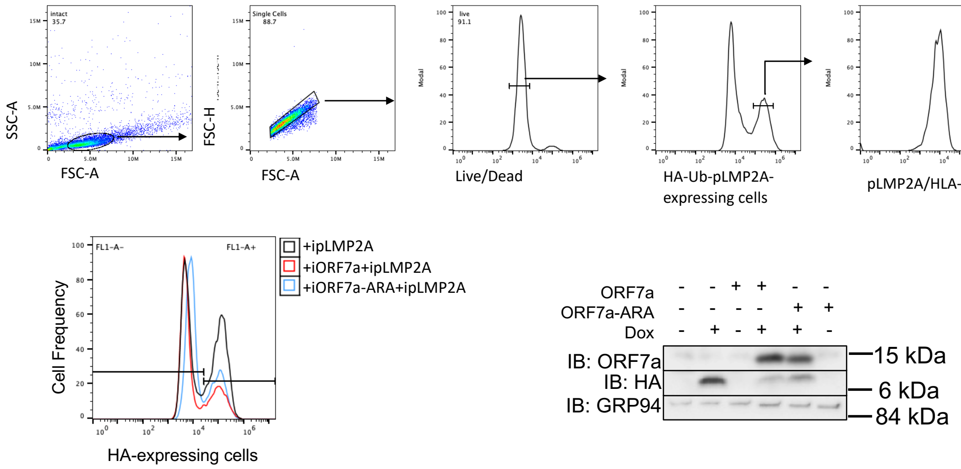


**Fig. S7**: Representative gating strategy for the FACS analysis of p-LMP2A-HLA-A2 complexes in the various cell lines (top). Proportion of cells expressing HA-Ub-pLMP2A fusion protein in induced cells and accompanying western blot analysis showing expression of ORF7a, HA-Ub-pLMP2A and GRP94 as loading control, in uninduced and induced cells (+Dox for 24 h, below).

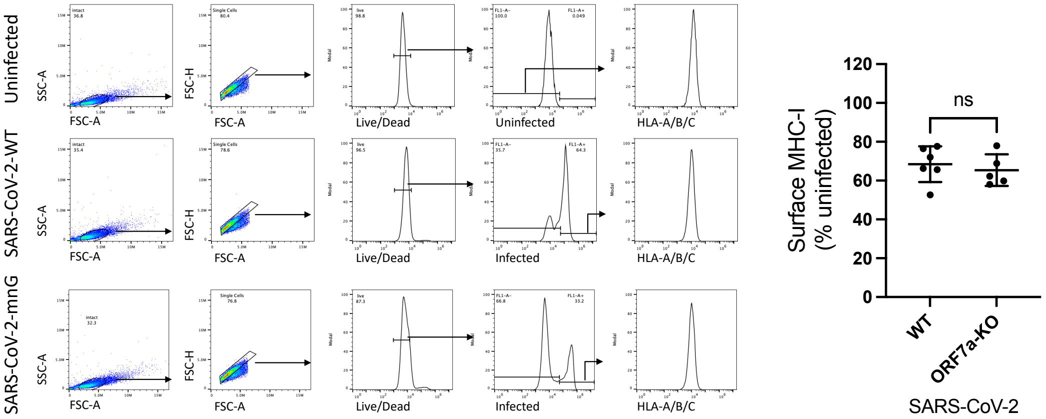


**Fig S8**: Infection with SARS-CoV-2 virus expressing or not expressing ORF7a does not alter MHC-I downregulation. Vero E6 or HEK293T-Ace2 cells were infected with SARS-CoV-2$_{WA1}$ or recombinant SARS-CoV-2 expressing fluorescent protein mNeoGreen (CoV-2-mNG, effectively an knockout of ORF7a) (42) (MOI = 10). 24 h post infection, cells were collected for flow cytometry analysis for surface MHC-I (n = 4). Cells infected the WT virus were identified by intracellular staining for spike protein and fluorescence from NeonGreen was used to identify cells infected with the ORF7a-null virus. Mean fluorescence intensity of MHC-I staining with W6/32 was normalized to uninfected group.

**Table S1.** Primers used in the generation of various plasmids used in the study

| Clone name | Primer name | Sequence (5' -3') |
|---|---|---|
| pCAGGS-ORF3a | ORF3a-f | GGCAAAGAATTCACCATGGATTTGTTTATGAGAATCTTCACAATTGGAACTGTAACTTTGAAGCAAGG |
| | NoTag-r | CATGGTGAATTCTTTGCCAAAATGATGAGACAGCACAATAACC |
| pCAGGS-ORF7a | ORF7a-f | GGCAAAGAATTCACCATGAAAATTATTCTTTTCTTGGCACTGATAACACTCGCTACTTG |
| | NoTag-r | CATGGTGAATTCTTTGCCAAAATGATGAGACAGCACAATAACC |
| pCAGGS-ORF7a-ARA | ARA-f | CACACTCGCAAGAGCGACAGAATGAGCTAGCAGATCTTTTTCCC |
| | ARF-R | TCATTCTGTCGCTCTTGCGAGTGTGAAGCAAAGTGTTATAAACACTATTGC |
| pCAGGS-ORF8 | ORF8-f | GGCAAAGAATTCACCATGAAATTTCTTGTTTTCTTAGGAATCATCACAACTGTAGCTGC |
| | ORF8-r | CATGGTGAATTCTTTGCCAAAATGATGAGACAGCACAATAACC |
| pTRIPZ-puro-ORF7a | IndORF7a-f | TAGTGAACCGTCAGATCGCACCGGTATGAAAATTATTCTTTTCTTGGC |
| | IndORF7a-r | GCCACGCCTCGAGACGCGTGTCATTCTGTCTTTCTTTTGAGTG |
| pTRIPZ-puro-ORF7a-ARA | IndORF7a-ARA-r | GCCACGCCTCGAGACGCGTGAATTCTCATTCTGTGGCTCTGGC |
| pTRIPZ-puro-HA-Ub | HAUb-f | TAGTGAACCGTCAGATCGCACCGGTATGTACCCATACGATGTTCCG |
| | HAUb-r | GCCACGCCTCGAGACGCGTGAATTCACCACCTCTCAGACG |
| pTRIPZ-puro-HA-Ub-pLMP2A | Ub-LMP2a-f | CCTGGTCCTGCGTCTGAGAGGTGGTAGCGGATCCTGTCTGGGCGGCCTGCTGACCATGGTGTAAGAATTCACGCGTCTCGAGGCGTGGCC |

**Table S2.** Primers used for qPCR analysis

| Gene name | Primer name | Sequence (3'-5') |
|---|---|---|
| MHC-I heavy chain | panHC-RT-f | GGGCTACGTGGACGACAC |
| | panHC-RT-r | CTCTGGTTGTAGTAGCCGCGC |
| ORF7a | ORF7a-RT-f | CTTCTGGAACATACGAGGG |
| | ORF7a-RT-r | CCTCTTGTCTGATGAACAG |
| GAPDH | GAPDH-RT-f | TCAAGGCTGAGAACGGGAAG |
| | GAPDH-RT-r | CGCCCCACTTGATTTTGGAG |
